# N-cadherin in osteolineage cells restrains breast cancer cell growth via inhibition of a PI3K-dependent, Tgf-β1-driven feed-forward loop

**DOI:** 10.1101/2025.10.22.683948

**Authors:** Toshifumi Sugatani, Kate Yeo, Marco Campioli, Hillary Fujimoto, Roberto Civitelli

## Abstract

Tumor growth and metastases are affected by interactions between tumor and microenvironment cells. We have reported the presence of *Sp7*-positive cells with an osteogenic signature in primary mouse breast cancer, where they stimulate tumor growth; and genetic ablation of *Cdh*2 (encoding N-cadherin) in these cells enhances their pro-tumorigenic action. To study the molecular mechanisms of this biologic system, we used MC3T3 cells, phenotypically similar to tumor associated osteolineage cells. Ablation of *Cdh2* in MC3T3 cells enhances PI3K-Akt-β-catenin signaling in response to transforming growth factor-β1 (Tgf-β1), resulting in increased production of Tgf-β1. Interference with PI3K activity is mediated by N-cadherin binding to PI3K components, p85α and p100, resulting in reduced activation of PI3K-Akt-β-catenin signaling. Downstream, *Cdh2* ablation enhances Tgf-β1-induced binding of Sp1 and Lef-1 to the *Tgfb1* promoter, leading to increased promoter activity and enhanced Tgf-β1 production. This is associated with miR-21 up-regulation and decreased expression of Pten, a PI3K inhibitor. MC3T3 cells promote growth of breast cancer cells (BCC) when co-cultured *in vitro* or co-injected in mouse mammary fat pad, and *Cdh2*-deficiency enhances this pro-tumorigenic effect. Notably, genetic ablation of the Tgf-β1 receptor subunit, *Tgfbr1,* in BCC abrogates the pro-tumorigenic action of MC3T3 cells and its enhancement by *Cdh2* ablation. Finally, Sp7-driven *Tgfbr1* ablation in mice also reduces the growth of BCC in the mammary fat pad. Thus, autocrine Tgf-β1 production via PI3K-Akt-β-catenin signaling is a key mechanism by which osteolineage cells promote BCC growth, an action restrained by Ncad via interference with PI3K components.

**Highlights:**

- In tumor-associated osteolineage cells, N-cadherin reduces the growth of breast tumors; it also inhibits Tgf-β1-activated PI3K/AKT/β-catenin signaling
- Tgf-β1 produced by tumor microenvironment cells stimulates breast cancer cell growth and further autocrine production of Tgf-β1
- The anti-tumorigenic action of N-cadherin is mediated by a braking effect on a Tgf-β1-driven, pro-tumorigenic, feed-forward cycle between microenvironment and tumor cells.

## 1. Introduction

Tumor progression is in part driven by interactions between tumor microenvironment (TME) and cancer cells (1). Expression of the calcium-dependent cell-cell adhesion molecule N-cadherin (Ncad), encoded by the *Cdh2* gene, by tumor cells is the hallmark of epithelial-to-mesenchymal transition, resulting in the acquisition of an aggressive tumor phenotype and increased metastatic potential (2). Accordingly, antibody neutralization of Ncad diminishes migration of cancer cells and reduces metastases in xenograft models (3). Ncad inhibitors can also suppress the growth of Ncad overexpressing pancreatic tumors (4) and improve response to chemotherapy in preclinical models (5). However, opposite effects of Ncad have also been described; for example, Ncad functions as a growth suppressor in K-ras-induced murine pancreatic tumors (6), and interference with Ncad promotes tumor growth in mouse melanoma (7). More to the point, loss of Ncad was associated with poorer prognosis in hepatocellular carcinomas (8), and clinical trials using Ncad antagonists for potential anti-tumorigenic action have been quite inconsistent (7,9,10), highlighting the need of a better understanding of the role of Ncad in tumorigenesis and cancer progression.

While Ncad in tumor cells has been extensively studied, less attention has been given to how Ncad in TME cells contributes to the interaction between the tumor and the tissue in which the cancer grows. In one study, heterotypic interactions between Ncad in cancer-associated fibroblasts (CAF) and E-cadherin (Ecad) in tumor cells favored tumor cell spreading (11). Likewise, interactions between Ncad in osteogenic cells and Ecad in tumor cells were proposed as a mechanism leading to formation of the pre-metastatic niche in bone (12). These findings suggest that Ncad in osteogenic or fibroblastic cells present in the TME may be pro-tumorigenic. However, contrary to such expectations, we have found that tumor cells inoculated in the mouse mammary fat pad grow larger in mice with *Cdh2* gene ablation targeted to osteolineage cells, with increased incidence of metastases (13). We also find that cells expressing the osteolineage factor, Osterix (Osx) encoded by the *Sp7* gene, are present in mouse breast tumors growing from inoculated breast cancer cells (BCC) (14), and that co-injection of Osx+ TME cells with BCC leads to enhanced tumor growth (14). Therefore, Ncad in these tumor-associated Osx+ cells is anti-tumorigenic, a function opposite to what reported for Ncad in CAF (11). Although they both produce collagen matrix, fibroblasts and osteoblasts differ in many ways. Indeed, our transcriptomic analysis showed that tumor associated Osx+ cells have a phenotypic signature more like osteogenic cells than CAF (13). Of note, the pro-tumorigenic action of Ncad in CAF was attributed to its cell-cell adhesion function that favors cells spreading (11). However, Ncad is also involved in cell signaling, and in bone cells it negatively regulates Wnt/β-catenin (15,16) and phosphoinositide 3-kinase (PI3K) (17). Accordingly, we found that PI3K, β-catenin, Ras, and extracellular signal-regulated kinase (Erk) pathways are all hyper-activated in tumor-associated Osx+ cells upon *Cdh2* ablation (13). Furthermore, gene promoters containing consensus sequences for specificity protein 1 (Sp1), lymphoid enhancer-binding factor-1 (Lef1), myc-associated zinc finger protein (Maz), and nuclear factor of activated T-cell (Nfat) transcription factors are all upregulated in *Cdh2*-deficient tumor associated Osx+ cells compared to WT cells (13).

To study the molecular mechanisms by which *Cdh2* ablation in Osx+ cells results in enhanced tumor growth, we generated an osteogenic cell line where *Cdh2* was ablated by CRISPR/Cas9 gene editing. We find that in *Cdh2* deficient osteogenic cells PI3K-Akt-β-catenin signaling is hyper-activated in response to transforming growth factor-β1 (Tgf-β1), resulting in enhanced Sp1 and β-catenin/Lef1 binding to the Tgfβ-1 promoter, transcriptional activity, and in turn, Tgf-β1 production, associated with enhanced growth of BCC. Importantly, both accentuated Tgf-β1 production and support of tumor growth brought about by *Cdh2* deletion were abrogated by ablation of *Tgfbr1*, a Tgf-β1 receptor subunit, in cancer cells, demonstrating that Ncad in osteolineage cells functions as a break to a Tgf-β1-driven positive feedback cycle that promote tumor growth through interference with PI3K-Akt-β-catenin signaling.

## 2. Material and Methods

### Antibodies, primers and other key reagents

The following antibodies were used: N-cadherin (1:1000, Cell Signaling), p-PI3K-p85α (Abcam, 1:500), PI3K-p85α (1:1000, Cell Signaling), p-Akt (Ser473) (1:2000, Cell Signaling), Akt (1:2000, Cell Signaling), p-β-catenin (Ser552) (1:500, Cell signaling), β-catenin (1:1000, Cell signaling), histone H3 (1:3000, Cell signaling), Pten (1:1000, Cell Signaling), Tgf-β receptor I (Abcam, 1:1000), α-tubulin (1:1000, Cell Signaling), Sp1 (1:100 Millipore Sigma), Lef1 (1:100 Millipore Sigma), RNA polymerase II (1:100 Abcam), or normal rabbit IgG (1:100 Cell signaling). All PCR primers were purchased from OriGene and are listed in Suppl. Table 1. The PI3K inhibitor, wortmannin was obtained from Cell Signaling.

### Cell culture

MC3T3, an osteoblast precursor cell-line derived from mouse calvaria (18), were purchased from ATCC. The cells were maintained in αMEM media without ascorbic acid (Thermo Scientific) including 10% FBS (Biowest) and penicillin/streptomycin (Thermo Scientific). The PyMT-BO1-GFP-firefly luciferase (Luc) (BO1) cell line, kindly provided by Dr. Katherine Welibaecher (Washington University), was derived from a primary tumor in MMTV-PyMT mice, and modified to stably express GFP and Luc reporters (19). The 4T1-GFP-Luc (4T1) murine mammary tumor cell line, which stably expresses Luc and GFP reporters, was generously provided by Dr. David Piwinica-Worms (MD Anderson Cancer Center, Houston, TX) (13). These cells were maintained in DMEM (Thermo Scientific) including 10% FBS (Biowest) and penicillin/streptomycin (Thermo Scientific).

### Generation of genetically engineered cell lines

To generate cell lines lacking either *Cdh2* or *Tgfbr1*, we used CRISPS/Cas9 gene editing technology. Targeting plasmids encoding for Cas9 and guide RNA for either *Cdh2* or *Tgfb1* were co-transfected with their respective homology-directed repair (HDR) plasmid (Santa Cruz) into either MC3T3 cells or BO1 and 4T1 cells, respectively, using FuGENE (Promega). These targeting plasmids (Santa Cruz) were designed to identify and cleave either exons 2-5 of *Cdh2*; or exons 3-5 of *Tgfbr1*. The resulting double-strand breaks are repaired by the specific HDR plasmids, which incorporate the puromycin resistance gene. Co-transfected cells were cultured for an additional 3 days in medium containing puromycin (8 μg/mL, Millipore Sigma), for selection of successfully transfected clones. Knockout of either gene was verified by verified by Western blotting of whole cell lysates (Sup. Fig. 1A and B).

### Generation of genetically engineered mice

Tet-off inducible *Sp7-Cre* (B6.Cg-Tg(Sp7-tTA,tetO-EGFP/cre)1Amc/J) mice and B6.Cg-Gt (ROSA) 26Sor Tm9(CAG-tdTomato)Hze/J (Ai9) mice were purchased from Jackson Laboratories. *Cdh2* floxed (*Cdh2^F/F^*) mice were obtained from Dr. Glenn L Radice (University of Pennsylvania School of Medicine, Philadelphia, PA). *Pten^F/F^* (B6.129S4-*Pten^tm1Hwu^*/J) and *Tgfbr1^F/F^* (*Tgfbr1^tm1.1Karl^*/KulJ) mice were purchased from Jackson Laboratories. Female mice were crossed with *Sp7-Cre* wild type male mice to generate *Cdh2^F/+^*, *Pten^F/+^* or *Tgfbr1^F/+^*, and conditionally deficient mice (*Cdh2^ΔSp7^*, *Pten ^ΔSp7^*, or *Tgfbr1 ^ΔSp7^*) were obtained by crossing *Cdh2^F/+^, Pten^F/+^* or *Tgfbr1^F/+^* with *Sp7-Cre* male mice and *Cdh2^F/F^*, *Pten^F/F^* or *Tgfbr1^F/F^* female mice, respectively. *Sp7-Cre* female mice were employed as a control mice. All animals were housed under pathogen-free conditions according to the guidelines of the Division of Comparative Medicine, Washington University School of Medicine. All animal protocols were approved by the Institutional Animal Care and Use Committee (Animal Welfare Assurance #D16-00245).

### Western blotting

After 6h serum starvation, MC3T3^WT^ or MC3T3^Cdh2KO^ cells were incubated with Tgf-β1 (20 ng/ml) for different times. Cells were harvested and whole-cell lysates prepared in RIPA buffer (Thermo Scientific). Nuclear extracts were obtained using the nuclear and cytoplasmic extraction reagent (Pierce). For all blots, 20 μg of proteins were loaded on the gel. Proteins were separated on 8–15 % SDS-PAGE and transferred to PVDF membrane (Millipore Sigma) by electrophoresis. Membranes were incubated with antibodies, as indicated in each figure, and bands were developed by chemiluminescence (Thermo Scientific). Band intensity was quantified by using ImageJ, a public domain program from the National Institutes of Health.

### Quantitative Real-Time PCR (qRT-PCR)

After 6h serum starvation, cell cultures were exposed to Tgf-β1 (20 ng/ml) (Pepro Tech) for 1h, and total RNA was isolated using the *mir*Vana™ miRNA isolation kit (Thermo Scientific). Clonal DNA was generated using the first strand cDNA synthesis system kit (OriGene). The amount of each gene product was determined using the OriGene qSTAR SYBR green kit in a 7300 Real-Time PCR System (Applied Biosystems). β_2_-microglobulin or U6 small RNA genes were used as controls. Quantification was obtained using the 2^-ΔΔCt^ method.

### Promoter Activity

Plasmids carrying promoter regions of *Sp1*, *Lef1*, or *Tgfb1* driving expression of Gaussia luciferase reporter gene (Gene Copoeia) were transiently transfected into MC3T3^WT^ or MC3T3^Cdh2KO^ cells. After 6-hour serum starvation, cells were incubated with Tgf-β1 (20 ng/ml) and harvested after 1h. Promoter activity was measured as bioluminescence intensity using the Pierce Gaussia Luciferase Glow Assay kit (Thermo Scientific).

### Immunoprecipitation (IP)

Whole-cell lysates of MC3T3 cells were prepared by sonication in RIPA buffer (Thermo Scientific), and the supernatant collected for IP assay. Part of the lysates were used as an input. Cell lysates were incubated with the antibody of interest overnight at 4°C. The immune complexes were pulled down using protein G or A magnetic beads, resuspended in 5X SDS sample buffer, and boiled at 100 °C for 5 min. After centrifugation, the supernatant was loaded on SDS-PAGE for immunoblot analysis.

### Chromatin immunoprecipitation (ChIP)

This was performed using the simpleChIP® enzymatic chromatin IP kit (Cell Signaling). In brief, after 6-h serum starvation and 1-h incubation with Tgf-β1 (20 ng/m), the cells were fixed with 37% formaldehyde to crosslink proteins to DNA, harvested and incubated with Micrococcal Nuclease to produce DNA/protein fragments. After incubation with antibodies of interest, the histone-protein immune complex was pulled down using Protein G or A magnetic beads. After reverse cross-linking, DNA was purified and amplified using PCR primers that detect the mouse *Tgfb1* promoter region (45) (ChIP-PCR). For quantitation of DNA-transcription factor binding (ChIP-qPCR), *Tgfb1* promoter primers were designed for optimization in the qSTAR SYBR green kit (OriGene). Quantitative binding was determined by calculating the amount of DNA obtained for each sample as a percentage of total DNA using the following formula: % of input = 2^(-ΔCt antibody)^ X 100, where Ct = cycle threshold, and ΔCt antibody = Ct total DNA – Ct antibody.

### In vitro tumor growth

BCC (BO1^WT^ and BO1^Tgfr1KO^; or 4T1^WT^ and 4T1^Tgfr1KO^) were seeded along with an equivalent number (2×10^5^ cells/well) of either MC3T3^WT^ or MC3T3^Cdh2KO^ cells and co-cultured in 24-well low-attachment plate (Fisher Scientific) with DMEM including 5% FBS and penicillin/streptomycin for 10 days. The medium was changed every 4 days. Luciferase intensity (Promega) was measured as an index of in vitro tumor growth

### In vivo tumor growth

For mammary fat pad inoculation, BCC (BO1^WT^ and BO1^Tgfr1KO^; or 4T1^WT^ and 4T1^Tgfr1KO^) and an equivalent number of either MC3T3^WT^ or MC3T3^Cdh2KO^ cells were mixed with growth factor-reduced matrigel (BD Biosciences) and co-injected into the mammary tissue of 8-week-old female C57BL/6 mice (Jackson Laboratories) under anesthesia with isofluraneMice were housed in standard temperature-and humidity-controlled environment with a 12h/12-h light/dark cycle. Tumor growth was monitored by measuring tumor volume (mm^3^) at each time-point using a caliper and calculated as: tumor volume = 0.5 X (minimum length)^2^ X (maximum length). At day 15, tumors were excised and immediately weighted. Animal procedures were approved by the Institutional Animal Care and Use Committee at Washington University (protocol number 20-0029) and followed the Animals in Research: Reporting of In Vivo Experiments (ARRIVE) guidelines.

### Statistical analysis

Unless otherwise indicated, group data are presented as boxplots with median (central bar) and interquartile range. Whiskers represent highest and lowest values. Statistical analyses were performed using Prism version 10 (GraphPad Software, Inc., La Jolla, CA, USA). Groups were compared by one-away analysis of variance (ANOVA), or two-way ANOVA to determine the contribution of multiple independent variables, as appropriate. In the latter case, Sidak multiple t-test was used for post hoc group comparisons.

### Data Availability

The data generated in the study are available upon request from the corresponding author.

## 3. Results

### Tgf-β1 hyperactivates non-canonical Tgf-β-signaling pathways in *Cdh2* ablated osteogenic cells

Osx+ cell present in the TME of breast cancer are rare (14) and in our hands have proven very difficult to grow in culture. Therefore, to study the molecular mechanisms by which Ncad in osteolineage cells exerts its anti-tumorigenic action and validate the bioinformatic analyses we previously reported (13), we turned to MC3T3 cells. These are undifferentiated but committed osteolineage cells (18) that when grown in osteogenic media undergo differentiation. They express *Osx* and other osteolineage factors (20); thus, they are phenotypically similar to Osx+ cells we found present in the TME of breast tumors growing from inoculated BCC. To test how cell signaling is affected by Ncad, we generated *Cdh2*-deficient MC3T3 cells (MC3T3^Cdh2KO^) using CRISPR/Cas9 gene editing. We monitored PI3K, Akt and β-catenin signaling in response to Tgf-β1 in MC3T3^Cdh2KO^ cells, since Tgf-β1 activates these signaling systems (21,22) and it is a known pro-tumorigenic factor involved in the paracrine “vicious cycle” between tumor and TME cells (23). Ncad protein abundance did not change within 60 minutes in response to Tgf-β1 in WT cells, and we confirmed that Ncad was undetectable in MC3T3^Cdh2KO^ cells (Fig.1A and Sup Fig.1A). While PI3K p85α and Akt protein abundance was not affected by Tgf-β1 exposure in both MC3T3^WT^ and MC3T3^Cdh2KO^ cells, accumulation of phospho-PI3K p85α and phospho-Akt (Ser473) rapidly increased upon exposure to Tgf-β1, and more abundantly so in MC3T3^Cdh2KO^ relative to MC3T3^WT^ cells (Fig.1A, Fig.2A and B). Furthermore, β-catenin phosphorylation at serine 552, which favors β-catenin accumulation in the nucleus and its transcriptional activity (24,25), was stimulated by Tgf-β1 more rapidly in whole cell lysates of MC3T3^Cdh2KO^ than in those of MC3T3^WT^ cells. Although 60 min after Tgf-β1 exposure, p-β-catenin (Ser552) accumulation was not substantially different between wild type or mutant cells (Fig. 1A and Fig. 2C), β-catenin was more abundant in the nuclear fraction of MC3T3^Cdh2KO^ than of MC3T3^WT^ cells (Fig. 1B and Fig. 2D), consistent with enhanced β-catenin nuclear translocation and transcriptional activity upon Tgf-β1 exposure in the mutant cells. Thus, Tgf-β1 favors β-catenin signaling via a non-canonical mechanism that is hyperactivated in Ncad-deficient osteogenic cells.

**Figure 1:**
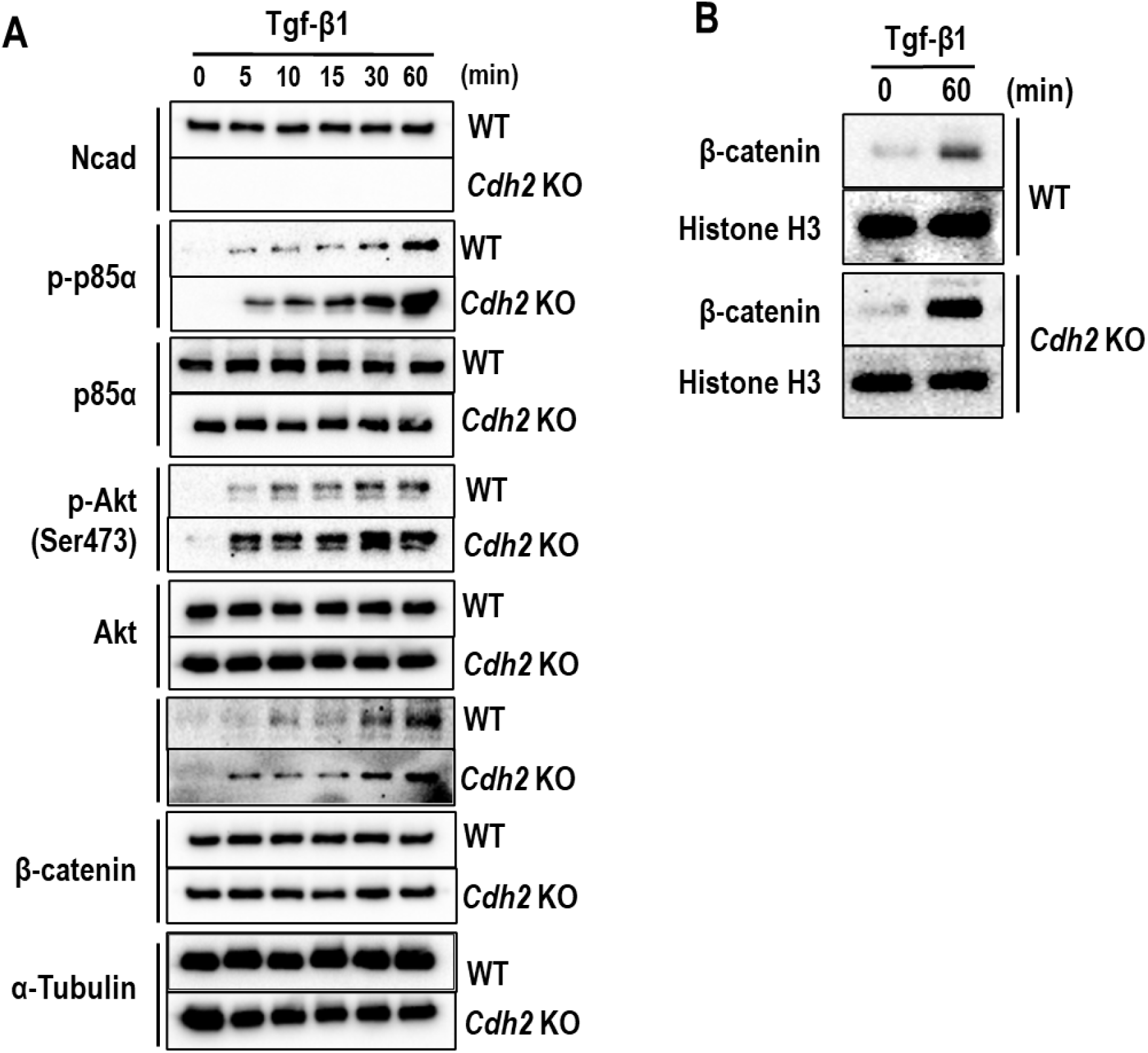
Hyperactivation of noncanonical Tgf-β1 signaling in *Cdh2* KO osteogenic cells. Confluent MC3T3^WT^ and MC3T3-E1*^Cdh2^*^KO^ cells were exposed to Tgf-β1 (20 ng/ml) for different times, as indicated. (A) Western blot of whole cell lysates using anti p-p85α, p85α, p-Akt (Ser473), Akt, Ncad, and p-β-catenin (Ser552) antibodies, with α-tubulin as loading control. (B) Nuclear extracts were obtained from confluent MC3T3^WT^ cultures for Western blot analysis using antibodies against total β-catenin and histone H3 as loading control.

**Figure 2:**
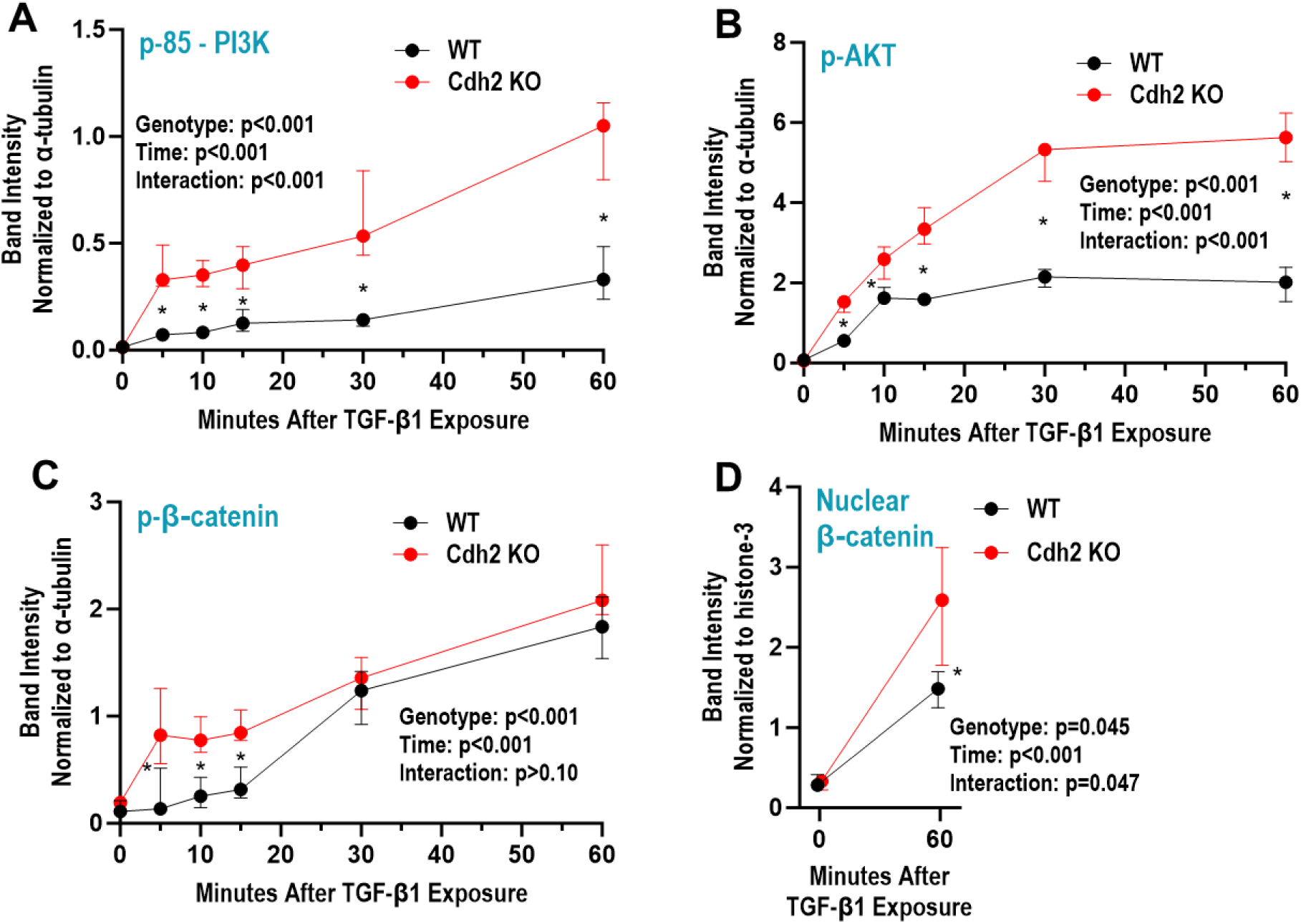
Quantitative analysis of noncanonical Tgf-β1 signaling in *Cdh2* KO osteogenic cells. The intensity of Western blot bands for (A) p-p85, (B) p-AKT, (C) p-β-catenin, and (D) nuclear β-catenin in MC3T3^WT^ and MC3T3-E1*^Cdh2^*^KO^ cells in response to Tgf-β1 (20 ng/ml) was quantified by using ImageJ, and normalized to bands for α-tubulin or histone-3, as indicated. Data are presented as median and range of triplicate experiments (Fig. 1 and Sup Fig. 1) and were analyzed by two-way ANOVA. Asterisks indicate p<0.05 for comparisons between the two cell lines at each time point (Sidak multiple comparison test).

### Increased expression of PI3K/Akt and β-catenin target genes in MC3T3^Cdh2KO^ cells

Consistent with enhanced signaling response to Tgf-β1, abundance of *Sp1* mRNA, downstream target of PI3K/Akt (26,27), and *Lef1* mRNA, a key mediator of the β-catenin signaling (28), was significantly higher after Tgf-β1 exposure in MC3T3^Cdh2KO^ cell than in MC3T3^WT^ cells (Fig. 3A and B). Moreover, Tgf-β1 induced *Tgfb1* mRNA expression to a larger extent in *Cdh2* ablated than in WT cells (Fig. 3C), suggesting the presence of an autocrine feed-forward regulation of Tgf-β1 in MC3T3^WT^ cells. Accentuated *Sp1*, *Lef1* and *Tgfb1* gene expression in *Cdh2*-deficient cells is most likely the consequence of increased transcriptional activity since promoter activities of all 3 genes was remarkably higher in MC3T3^Cdh2KO^ cells than in MC3T3^WT^ cells in response to Tgf-β1 (Fig. 3D-F). Based on these results, we hypothesize that Tgf-β1 in osteolineage cells up-regulates its own promoter *via* activation of PI3K/Akt and β-catenin and binding of Sp1 and Lef1 to their cognate sequences, a mechanism enhanced in *Cdh2*-deficient cells. Indeed, PCR analysis following chromatin immunoprecipitation revealed Sp1 and Lef1 reactive bands in nuclear extracts of MC3T3^WT^ cells, only after Tgf-β1 exposure (Fig. 3G), demonstrating that Tgf-β1 recruits Sp1 and Lef1 to the *Tgfb1* promoter. Importantly, Sp1 and Lef1 binding activities on the *Tgfb1* promoter were significantly higher in MC3T3^Cdh2KO^ cells than in MC3T3^WT^ cells in response to Tgf-β1 (Fig. 3H and I), indicating that PI3K-Akt and β*-*catenin hyperactivation by Tgf-β1 in the absence of Ncad results in enhanced Sp1 and Lef1 transcriptional activity to up-regulate autocrine *Tgfb1* expression.

**Figure 3:**
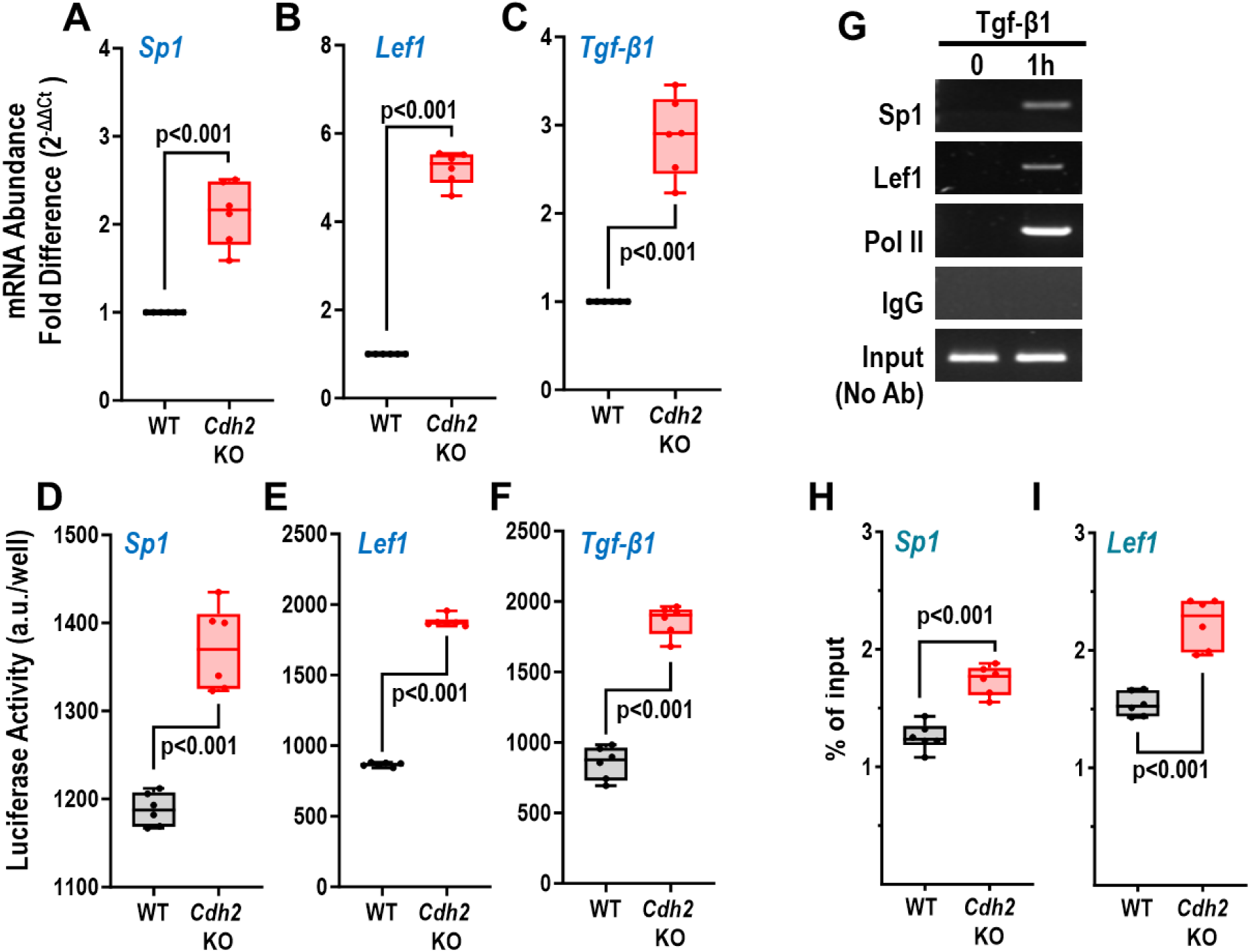
Enhanced *Sp1*, *Lef1*, and *Tgfb1* gene expression, transcriptional activity, and Sp1, Lef1 occupancy of the *Tgfb1* promoter in *Cdh2* KO osteogenic cells. Confluent MC3T3^WT^ and MC3T3-E1*^Cdh2^*^KO^ cells were exposed to Tgf-β1 (20 ng/ml) for 1 hour. Quantitative RT-PCR (qPCR) analysis of (A) *Sp1*, (B) *Lef1*, and (C) *Tgfb1* mRNA; (D) *Sp1*, (E) *Lef1*, and (F) *Tgfb1* promoter-firefly luciferase activity in WT and mutant cells. (G) *Tgfb1* promoter PCR products of MC3T3^WT^ chromatin immunoprecipitates obtained using Sp1, Lef1, polymerase II (Pol II), or anti-IgG antibodies, or on input sample (no immunoprecipitation) after 1h exposure to Tgf-β1. Abundance of Sp1 (H) or Lef1 (I) present on the *Tgfb1* promoter determined by qRT-PCR of the *Tgfb1* promoter. Groups were compared by one-way ANOVA.

We then asked whether phosphorylation of β*-*catenin at Ser552 is via the PI3K/Akt system. MC3T3^WT^ cells were incubated with or without the PI3K inhibitor, wortmannin, before exposure to Tgf-β1. Western blot analysis showed that wortmannin completely prevented Tgf-β1-induced Akt phosphorylation and reduced β*-*catenin phosphorylation at Ser552 (Fig. 4A). Accordingly, *Sp1* and *Lef1* mRNA expression was significantly reduced by the PI3K inhibitor (Fig. 4B). Of note, wortmannin did not inhibit Tgf-β1 induction of Smad2 phosphorylation in MC3T3^WT^ (Sup. Fig. 3), suggesting that PI3K-Akt signaling is activated independently of canonical Smad signaling by Tgf-β1 in osteolineage cells. Thus, β*-*catenin Ser552 phosphorylation by Tgf-β1 occurs predominantly via PI3K-Akt signaling in osteolineage cells.

**Figure 4:**
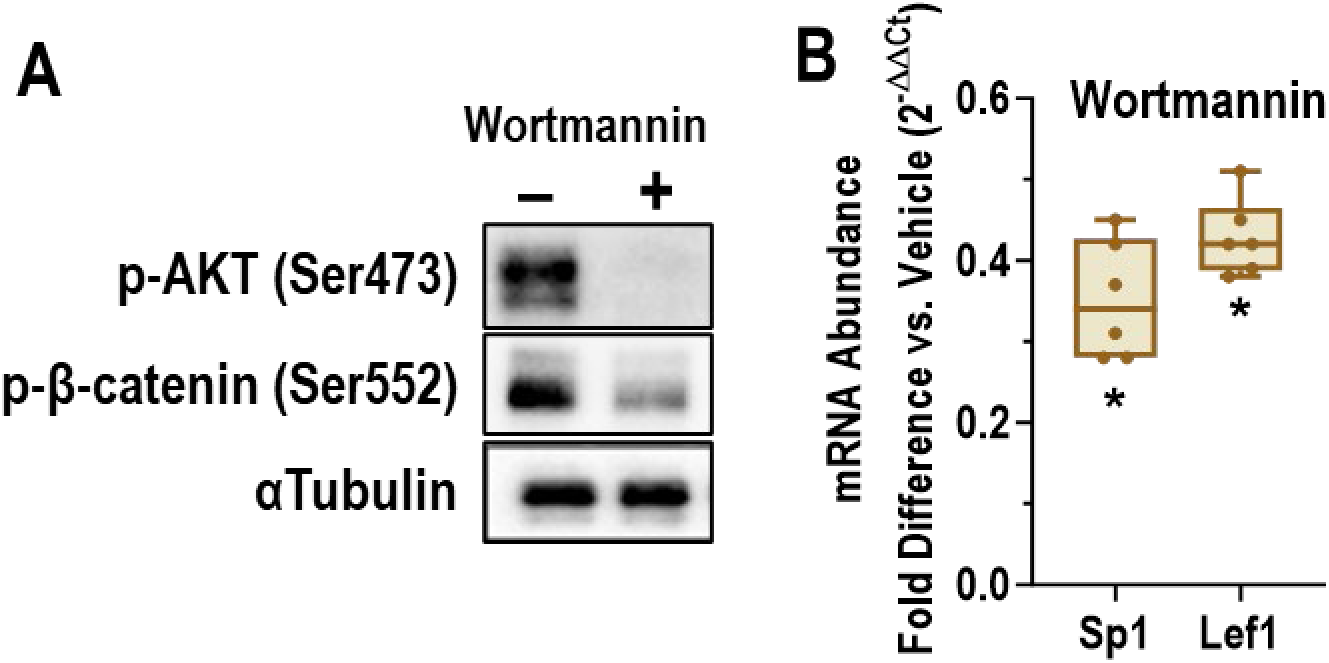
β-catenin Ser552 phosphorylation is sensitive to PI3K-Akt inhibition. Confluent MC3T3^WT^ cells were switched to αMEM without FBS and antibiotics for 6h, then incubated with the PI3K inhibitor, wortmannin (1 μM) or vehicle for 1 hour, followed by exposure to Tgf-β1 (20 ng/ml) for 1 hour. (A) Western blot analysis of whole cell lysates incubated with anti-p-Akt (Ser473), anti-p-β-catenin (Ser552), or anti-α-tubulin antibodies. (B) Quantitative RT-PCR analysis of *Sp1* and *Lef1* mRNA. Groups were compared by one-way ANOVA (\**p<0.001 vs*. vehicle).

We and others have shown that Ncad in osteolineage cells interferes with canonical Wnt signaling by binding to and sequestering Wnt components away from active signaling (29,30). To investigate whether a similar mechanism may be at play in hyperactivation of PI3K-Akt-β-catenin signaling in *Cdh2*-deficient cells in response to Tgf-β1, we tested binding of Ncad to PI3K components. Western blot analysis following immunoprecipitation demonstrated that Ncad-specific antibody pulled down p85α, and vice versa, a p85α-specific antibody pulled down Ncad in unstimulated MC3T3 cells (Fig. 5A and B). A p110-specific antibody also pulled down Ncad protein in the same cells (Fig. 5C), demonstrating that both PI3K components, p85α and p110 bind to Ncad. Phosphatase and tensin homolog (Pten) is a negative regulator of PI3K (31), and microRNA-21 (*miR-21*) targets *Pten* mRNA and interferes with its translation (32). We found that Tgf-β1 up-regulates *miR-21* expression in MC3T3^Cdh2KO^ relative to MC3T3^WT^ cells (Fig. 5D), and down-regulates Pten protein expression to a larger extent in MC3T3^Cdh2KO^ cells than in MC3T3^WT^ cells (Fig. 5E). Therefore, enhanced Tgf-β1-induced *miR-21* up-regulation and Pten down-regulation contribute to hyper-activation of Akt and β*-*catenin signaling in *Cdh2* deficient osteolineage cells.

**Figure 5:**
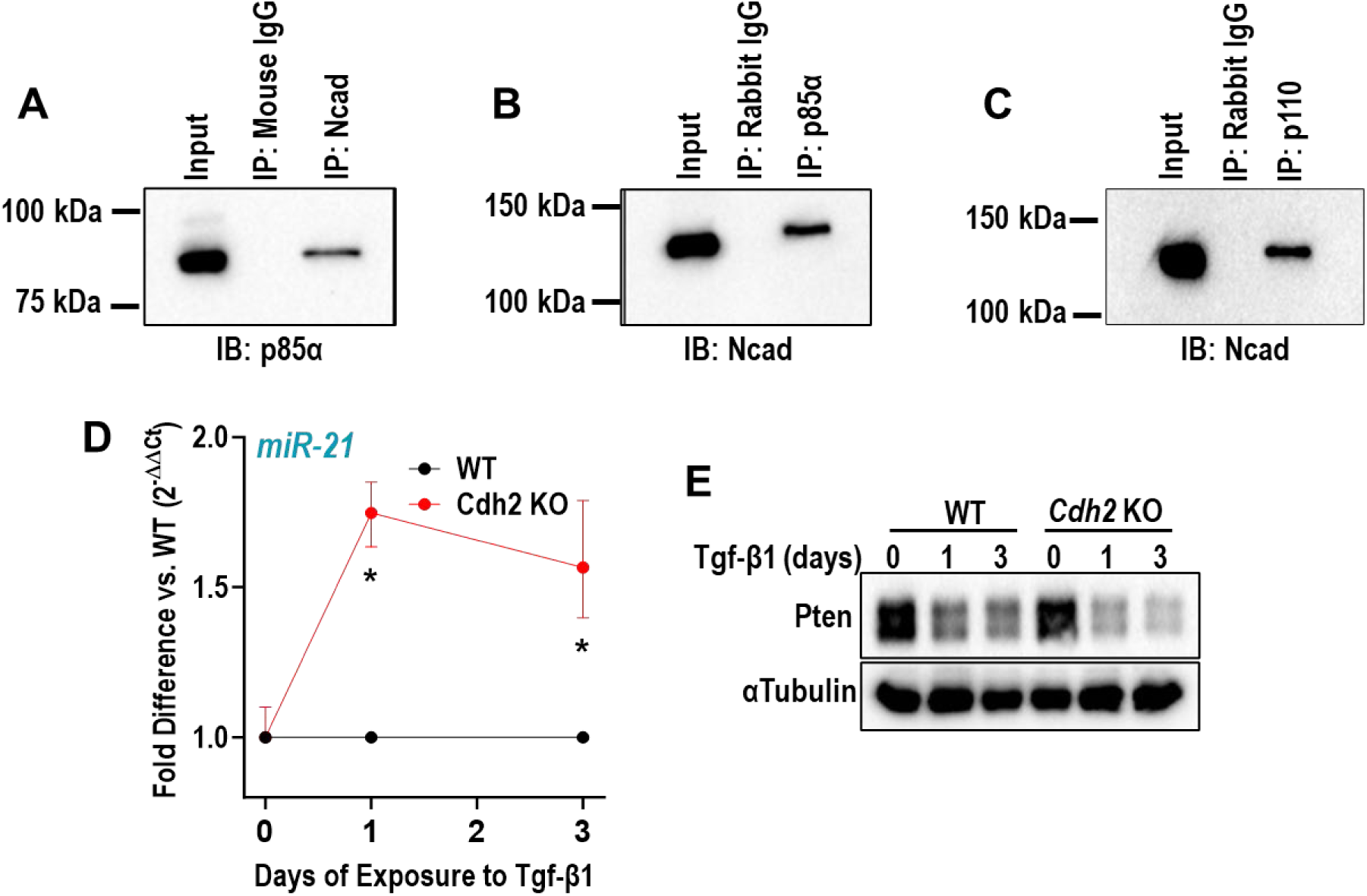
Ncad physically interacts with PI3K components and interferes with Pten and microRNA-21 expression. (A-C) Whole cell lysates of MC3T3 cells were immunoprecipitated using the antibodies indicated above the bands (IP) and immunoblotted using the antibodies indicated below the bands (IB). Input is the total protein lysate. (D) Confluent WT and *Cdh2* KO MC3T3-E1 cells were exposed to Tgf-β1 (20 ng/ml), and *miR-21* expression was determined by qPCR 1 and 3 days after Tgf-β1 exposure. Data represent median and range of 4 independent experiments. Groups were compared by one-way ANOVA (*p<0.001 vs. WT). (E) Whole cell lysates from similar cultures were used in Western analysis of Pten abundance in response to Tgf-β1; αTubulin was used as a loading control.

### Ncad in osteolineage cells is anti-tumorigenic in vitro and in vivo

The data shown thus far suggest that enhanced autocrine production of Tgf-β1 may be a mechanism by which *Cdh2*-deficient osteolineage cells are more pro-tumorigenic than wild type cells. To test this hypothesis, we generated *Tgfbr1*-deficient BO1 (BO1^Tgfr1KO^) and 4T1 (4T1^Tgfr1KO^) breast tumor cells, using CRISPR/Cas-9 technology. We targeted the Tgf-β receptor I subunit because it is necessary for activating downstream signaling upon Tgf-β1 ligation (33). When grown alone, BO1^WT^ or 4T1^WT^ cells form tumor spheroid aggregates whose volume can be assessed by bioluminescence, as these cells stably express firefly luciferase. When grown in co-culture with MC3T3^WT^ cells, tumor spheroid aggregates were more numerous and larger, and they were significantly larger still when BO1^WT^ or 4T1^WT^ cells were co-cultured with MC3T3^Cdh2KO^ cells (Fig. 6A-D and Fig. 7A-D). By contrast, BO1^Tgfr1KO^ or 4T1^Tgfr1KO^ cells formed significantly smaller and lesser spheroid aggregates when cultured alone relative to BO1^WT^ or 4T1^WT^ cells, suggesting that Tgf-β1 contained in the culture medium is an important stimulator of BO1^WT^ or 4T1^WT^ cell growth. More to the point, co-culture of BO1^Tgfr1KO^ or 4T1^Tgfr1KO^ cells with either MC3T3^WT^ or MC3T3^Cdh2KO^ cells failed to promote tumor spheroid growth (Fig. 6A, E-G and Fig. 7A, E-G), demonstrating that Tgf-β1 is a key factor mediating both the pro-tumorigenic action of osteolineage cells and the tumor enhancing effect of *Cdh2* deficiency.

**Figure 6:**
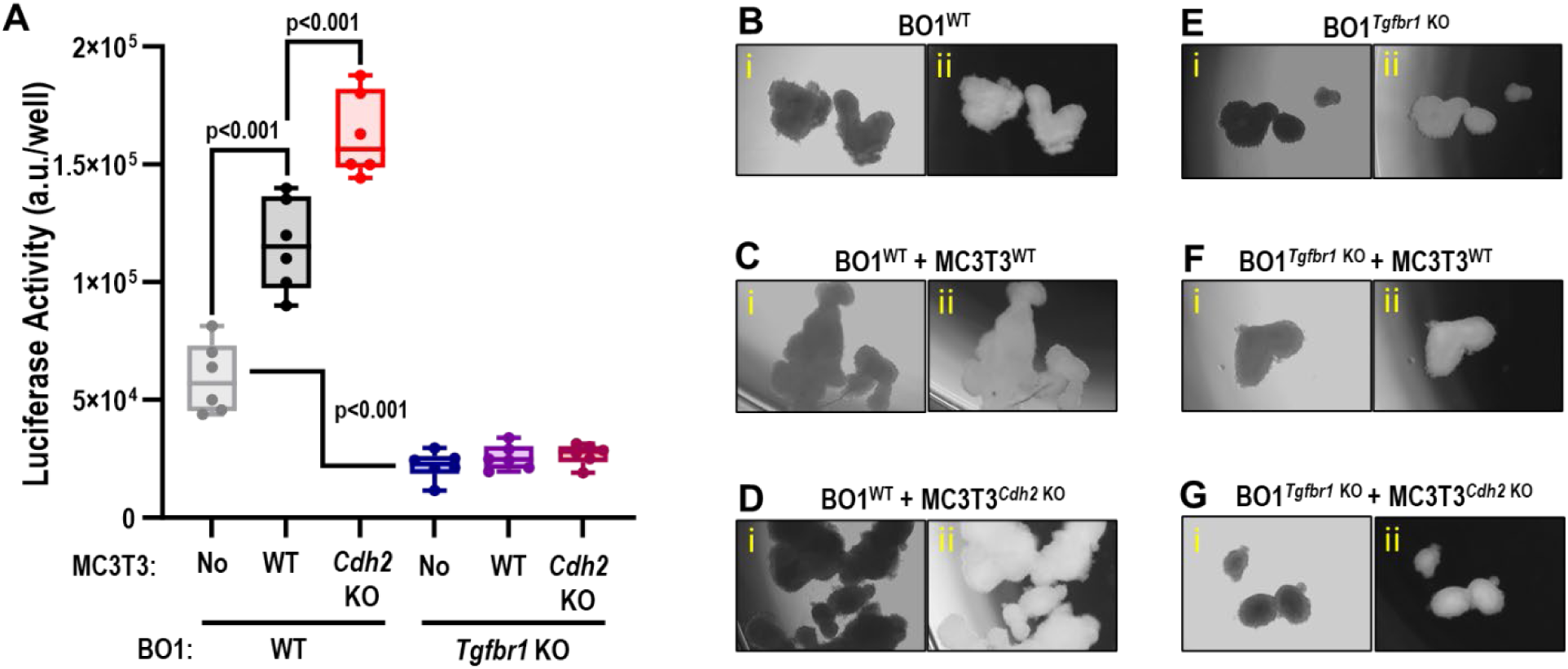
Lack of Ncad in osteogenic cells enhances BO1 cell growth in a Tgf-β receptor-dependent fashion *in vitro*. BO1^WT^ and BO1^Tgfr1KO^ breast cancer cells were cultured alone or in combination with either MC3T3^WT^ or MC3T3-E1*^Cdh2^*^KO^ cells for 10 days to form tumoral spheroids. (A) BO1 cell luciferase activity was determined as a measure of spherule growth after 10 days of co-culture. Groups were compared by one-way ANOVA. (B) Representative images of GFP-positive spheroids and aggregates of BO1^WT^ cells grown alone, (C) co-cultured with MC3T3^WT^, or (D) co-cultured with MC3T3-E1*^Cdh2^*^KO^ cells for 10 days. (E) GFP-positive BO1^tgfbr1KO^ spheroids and aggregates grown alone, (F) co-cultured with MC3T3^WT^, or (G) co-cultured with MC3T3-E1*^Cdh2^*^KO^ cells for 10 days. i) Brightfield, ii) fluorescence.

**Figure 7:**
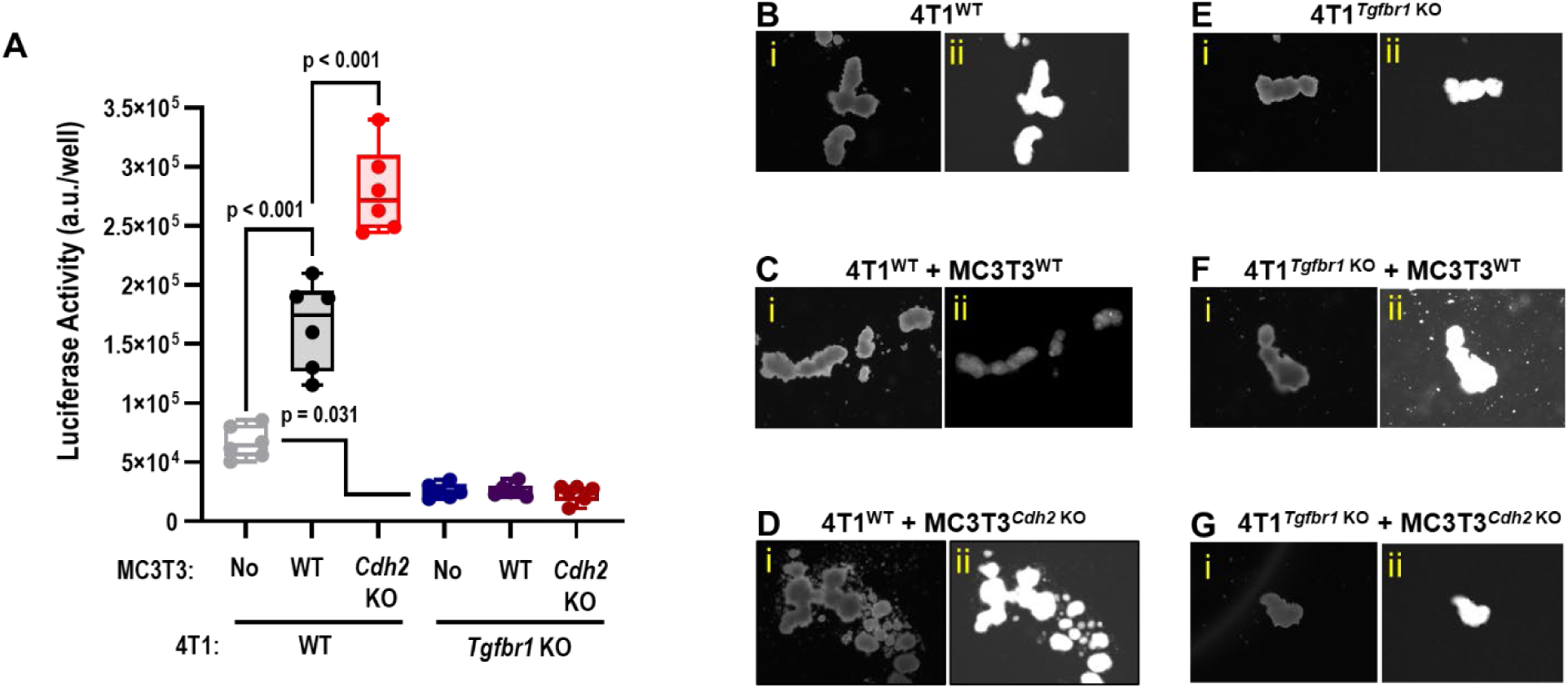
Lack of Ncad in osteogenic cells enhances 4T1 cell growth in a Tgf-β receptor-dependent fashion *in vitro*. 4T1^WT^ and 4T1^Tgfr1KO^ breast cancer cells were cultured alone or in combination with either MC3T3^WT^ or MC3T3-E1*^Cdh2^*^KO^ cells for 10 days to form tumoral spherules. (A) 4T1 cell luciferase activity was determined as a measure of spherule growth after 10 days of co-culture. Groups were compared by one-way ANOVA. (B) Representative image of GFP-positive 4T1^WT^ cells, (C) co-cultured with MC3T3^WT^, or (D) MC3T3-E1*^Cdh2^*^KO^ cells taken at day 10 of co-culture. (E) Representative image of GFP-positive 4T1^tgfbr1KO^ cells, (F) co-cultured with MC3T3^WT^, or (G) MC3T3-E1*^Cdh2^*^KO^ cells taken at day 10 of co-culture. i) Brightfield, ii) fluorescence.

To verify in vivo that *Cdh2* ablation in osteogenic cells stimulates tumor growth because of accentuated *Tgf-β1* expression, we co-inoculated genetically engineered BCC and osteogenic cells into the mammary fat pad of C57BL/6 female mice and monitored tumor volume every two days until tumor excision at day 15. Consistent with the in vitro results, co-inoculation of MC3T3^WT^ with BO1^WT^ or 4T1^WT^ cells resulted in significantly larger tumors compared to inoculation of BO1^WT^ or 4T1^WT^ cells alone, and co-injection of MC3T3^Cdh2KO^ cells further enhanced the growth of BO1^WT^ or 4T11^WT^ tumors (Fig. 8A-C and Fig. 9A-C). Unlike the *in vitro* results, however, there was no size difference between tumors growing from either WT or *Tgfbr1*-ablated BO1 or 4T1 cells, suggesting that pro-tumorigenic factors other than Tgf-β1 are present in the TME *in vivo* (Fig. 8A-C and Fig. 9A-C). Importantly, neither MC3T3^WT^ nor MC3T3^Cdh2KO^ cells supported the *in vivo* growth of tumors from BO1^Tgfr1KO^ or 4T1^Tgfr1KO^ cells (Fig. 8A-C and Fig. 9A-C), strongly suggesting that Tgf-β1 is a key pro-tumorigenic factor produced within the TME, and that the anti-tumorigenic effect of Ncad in osteolineage cells is primarily mediated by regulation of Tgf-β1 production.

**Figure 8:**
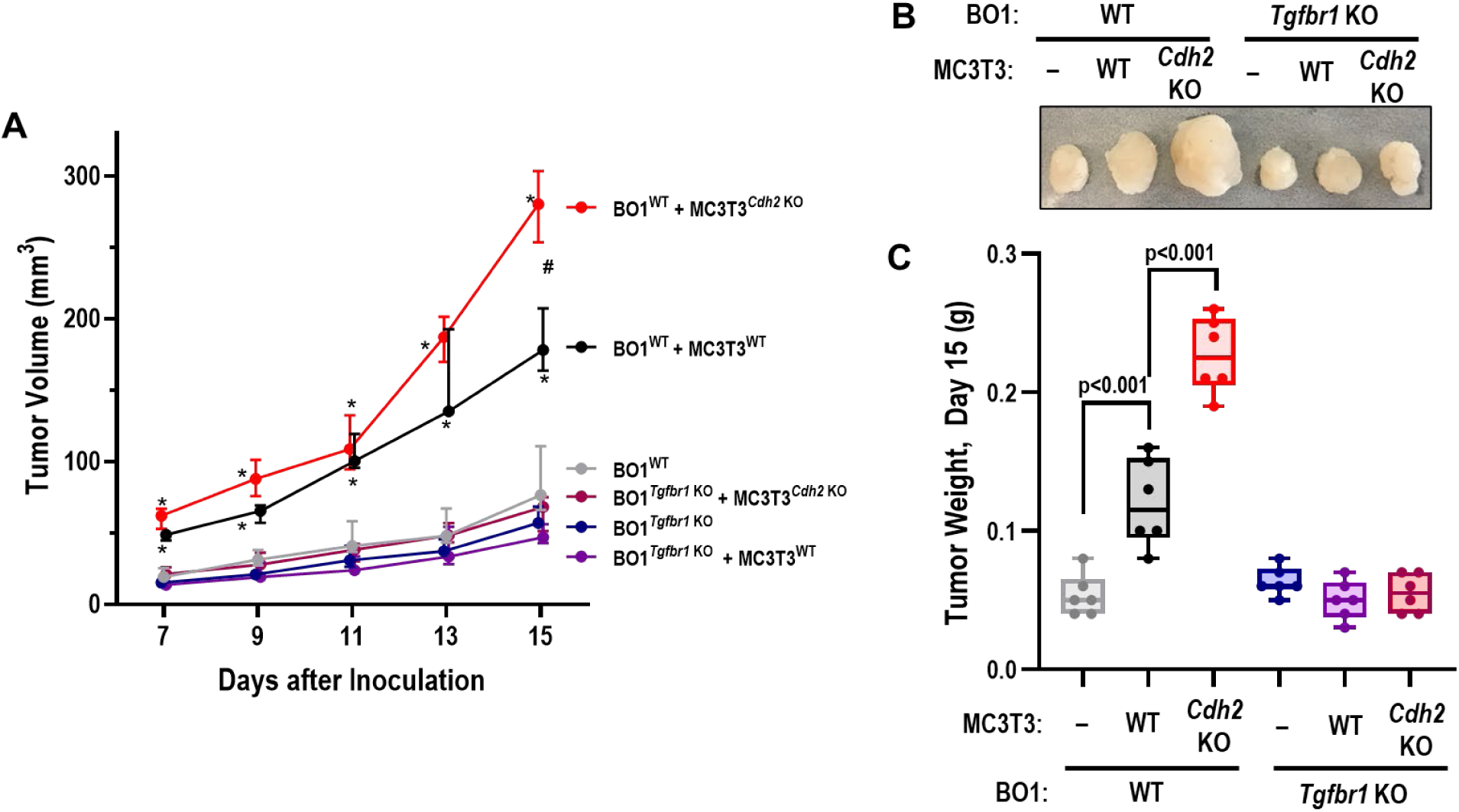
Lack of Ncad in osteogenic cells enhances BO1 cell growth in a Tgf-β receptor-dependent fashion *in vivo*. BO1^WT^ and BO1^Tgfr1KO^ breast cancer cells were inoculated into the mammary fad pad of female mice alone or with either MC3T3^WT^ or MC3T3-E1*^Cdh2^*^KO^ cells. (A) The volume of tumors developing from these inoculates was monitored by caliper from day 7 to 15. Data represent median and interquartile range. There was a significant interaction between time and cell type (p<0.001; two-way ANOVA); *p<0.01 *vs*. BO1^WT^; #p=0.007 *vs*. BO1^WT^ + MC3T3^WT^ (Tukey test for multiple comparisons). (B) Representative image of explanted tumors at day 15 after cell inoculation. (C) Weight of tumors explanted. Groups were compared by one-way ANOVA.

**Figure 9:**
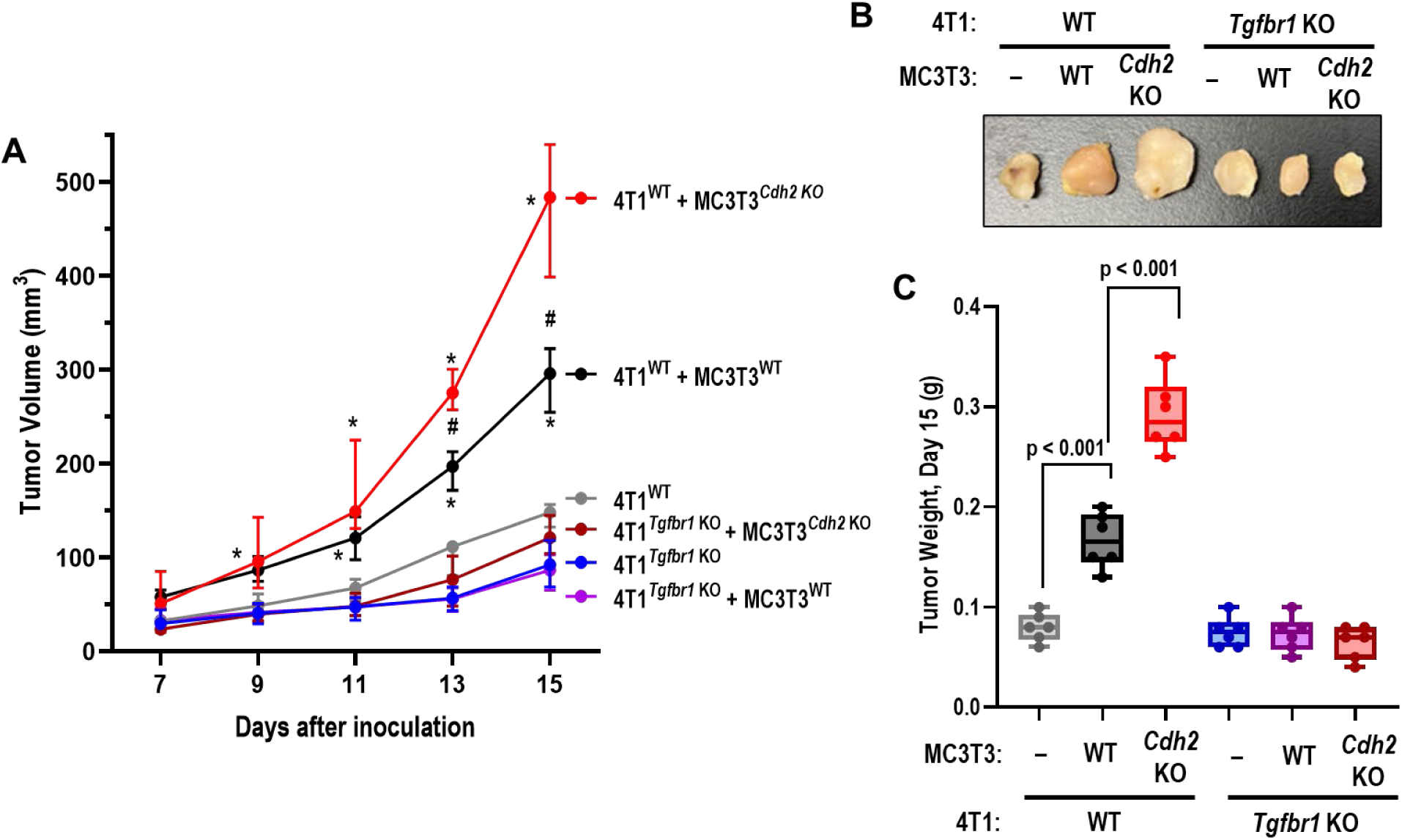
Lack of Ncad in osteogenic cells enhances 4T1 cell growth in a Tgf-β receptor-dependent fashion *in vivo*. 4T1^WT^ and 4T1^Tgfr1KO^ breast cancer cells were inoculated into the mammary fad pad of female mice alone or with either MC3T3^WT^ or MC3T3-E1*^Cdh2^*^KO^ cells. (A) The volume of tumors developing from these inoculates was monitored by caliper from day 7 to 15. Data represent median and interquartile range. There was a significant interaction between time and cell type (p<0.001; two-way ANOVA); *p<0.01 vs. 4T1^WT^; # p=0.01 vs. 4T1^WT^ + MC3T3^WT^ (Tukey test for multiple comparisons). (B) Weight of tumors explanted at day 15 after cell inoculation. Groups were compared by one-way ANOVA. (C) Representative image of explanted tumors.

To validate in vivo the role of Tgf-β1 produced by osteolineage cells in BCC growth and Ncad interference, we used Sp7-Cre (also called Osx-Cre) to generate mice with conditional ablation of *Cdh2* (*Cdh2 ^ΔSp7^*), as previously reported (34), or *Tgfbr1* (*Tgfbr1 ^ΔSp7^*), or *Pten* (*Pten ^ΔSp7^*). Inoculation of 10^5^ BO1 cells in the MFP resulted in larger tumors in *Cdh2 ^ΔSp7^* and *Pten ^ΔSp7^* mice compared to control, Sp7-Cre mice. Conversely, tumor growth was significantly slower between days 7 and 15 post-injection in *Tgfbr1 ^ΔSp7^* (Fig. 10 A). Accordingly, tumors excised at day 15 were significantly larger in *Cdh2 ^ΔSp7^* and *Pten ^ΔSp7^* mice relative to Sp7-Cre mice, whereas they were significantly smaller in *Tgfbr1 ^ΔSp7^* mice (Fig. 10B).

**Figure 10:**
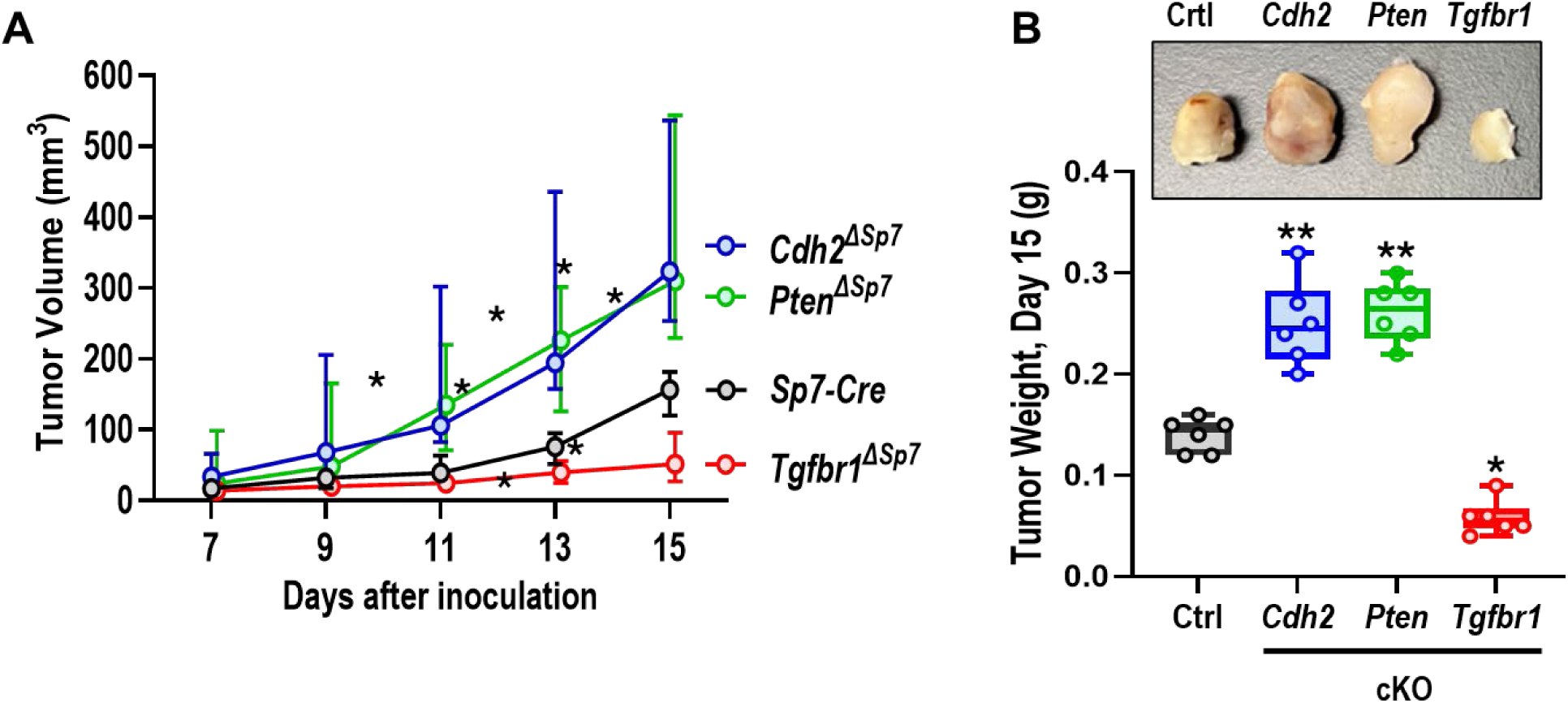
The non-canonical Tgf-β1 signaling in *Sp7*+ tumor stromal cells is involved in breast cancer growth. BO1 breast cancer cells were inoculated into the mammary fad pad of *Cdh2*, *Pten*, or *Tgfbr1* cKO female mice and their control (*Sp7-Cre*;Ai9) female mice. (A) The volume of tumors developing from these inoculates was monitored by caliper from day 7 to 15. Data represent median and interquartile range. There was a significant interaction between time and cell type (p<0.001; two-way ANOVA); *p=0.0165, **p=0.009, ***p<0.0001 vs. Sp7-Cre;Ai9. (B) Representative image of explanted tumors and weight of tumors explanted at day 15 after cell inoculation. Groups were compared by one-way ANOVA; *p=0.0002, **p<0.0001 vs. Ctrl.

## 4. Discussion

We discovered a new mechanism by which Ncad negatively modulates a Tgf-β1-driven positive feedback cycle through interference with PI3K-Akt-β-catenin signaling and transcriptional down-regulation of the *Tgfb1* promoter. In the context of the TME, where cells with an osteogenic signature are present, Ncad antagonizes the pro-tumorigenic effect of such Tgf-β1 feed forward, “vicious” cycle, thus reducing tumor growth.

Our previous work demonstrated the presence of pro-tumorigenic Osx+ cells in mice breast tumors, and *Cdh2* ablation in Osx-targeted cells resulted in enhanced growth of tumors implanted in the mammary fat pad (13,14). In this work, we studied the molecular mechanisms attending to the enhanced tumorigenic effect caused by lack on Ncad in Osx+ cells. We demonstrate that a cell line, MC3T3, which is phenotypically close to tumor-associated Osx+ cells (13), reproduce the pro-tumorigenic action of tumor associated Osx+ cells when co-cultured with breast cancer cells and when co-injected with breast cancer cells in the mouse mammary fat pad. We also show that such pro-tumorigenic action is enhanced when *Cdh2* is genetically inactivated in Osx+ cells, in vitro and in vivo, fully corroborating our previous results and validating the cell models we used for this molecular analysis. Ncad interference of the Tgf-β1-driven paracrine cross-talk between osteolineage microenviroment and tumor cells has important ramifications for both primary breast tumor progression, and bone metastasis, where interactions with resident bone cells is a key component of tumor cell colonization and growth (23).

Cadherins are integral components of the adherens junction, where they interact with α-, β-, and γ-catenin (35,36). Since β-catenin is also a key component of the Wnt signaling system, cadherins can also modulate Wnt/β-catenin signaling (37,38). Work from our laboratory and others demonstrated that cadherins interfere with signaling pathways that are key to osteogenic differentiation and skeletal development (15,39). Indeed, we have shown that ablation of *Cdh2* in osteogenic cells in mice reduces the number of bone marrow osteoprogenitors leading to low bone mass (34), and demonstrated that Ncad restrains parathyroid hormone bone anabolic effect and activation of β-catenin signaling via interaction with low-density lipoprotein receptor-related protein 6 (Lrp6) and parathyroid hormone receptor 1 in osteogenic cells (30). Others have shown that Ncad/Lrp6 interaction leads to β-catenin degradation and decreased Wnt signaling (29,40). Here, we find that Ncad functionally interacts with PI3K components, p85α and p110, suggesting that, in analogy to what occurs with Lrp5/6, Ncad may sequester p85α and p110 away from signaling. Therefore, in the absence of Ncad more p85α and p110 are available for activation through the Tgf-β/BMP receptor system. Indeed, overexpression or siRNA interference with *Cdh2* decreases or increases PI3K signaling, respectively (17), fully consistent with our present findings.

Downstream of PI3K activation by Tgf-β1, AKT phosphorylation results in “non-canonical” stabilization of β-catenin via direct phosphorylation in Ser552. Accordingly, both *Sp1* and *Lef1*, a Wnt signaling component, are transcriptionally up-regulated, and in turn bind to the *Tgfb1* promoter, resulting in transcriptional activation and Tgf-β1 production. These results are fully consistent with our previous RNA sequencing data, indicating that gene promoters containing Sp1 and Lef1 consensus sequences are up-regulated in Ncad-deficient tumor-associated Osx+ cells (13). We also found that Tgf-β1 downregulates Pten protein, an inhibitor of PI3K (31), most likely via Sp1-dependent up-regulation of miR-21 expression (27). Lack of Ncad does not alter basal Pten expression but accentuates Tgf-β1-dependent Pten downregulation and miR-21 up-regulation, reflecting Ncad action as a break to Tgf-β1-dependent PI3K activation. Of note, Pten has a tumor suppressive action (41,42). Indeed, we find that conditional ablation of *Pten* in Osx+ cells phenocopies enhanced tumor growth seen with *Cdh2* conditional ablation. Importantly, both the pro-tumorigenic effect of MC3T3 cells and its enhancement in the absence of Ncad are heavily dependent on Tgf-β/BMP signaling, as genetic ablation of *Tgfb1*, a key receptor subunit for Tgf-β/BMP signaling in tumor cells abrogated both effects in cell co-cultures and *in vivo*. Likewise, conditional *Tgfbr1* ablation in Osx+ cells also reduced BCC growth in vivo, demonstrating the key role of the Tgf-β/BMP system in paracrine modulation of tumor growth in the breast. However, other mechanisms may exist; the slower but equal growth of WT or*Tgfbr1* ablated BO1 cells when injected in the mouse suggests that some factors present in the TME stimulate cancer cell growth independently of Tgf-β/BMP signaling.

Our data reveal the existence of a pro-tumorigenic, feed-forward loop driven by Tgf-β1 (Fig. 11). We propose that tumor-associated osteogenic cells produce and respond to Tgf-β1, as osteoblasts do in normal bone remodeling. In the TME, Tgf-β1 secreted by osteolineage or other TME cells directly stimulates cancer cell proliferation and tumor growth, and feeds back onto the osteogenic cells to further stimulate its own synthesis. Mechanistically, Tgf-β1 receptor binding causes phosphorylation of the two main PI3K components, p85 and p100, and consequent Akt phosphorylation. In turn, activated Akt triggers downstream events all leading to increased transcriptional activity of the *Tgfb1* promoter: 1) Sp1 recruitment to the promoter; 2) β-catenin stabilization by Ser552 phosphorylation, nuclear translocation, and increased promoter occupancy, most likely by formation of β-catenin/Lef1 complexes; 3) increased miR-21 expression and post-transcriptional downregulation of the PI3K inhibitor, Pten. These events generate a positive feed-back loop that induces further autocrine Tgf-β1 production by TME osteolineage cells, fueling tumor growth. Ncad provides a break to this feed-forward “vicious” loop, by retaining the PI3K components, p85α and p110, away for signaling and reducing PI3K signaling and Tgf-β1 production. Therapeutically, targeting Ncad with inhibitors or neutralizing antibodies has not been very successful for oncologic indications (7,9,10), likely because of the multiple and sometimes opposing functions of Ncad in different cell types. For example, Ncad in CAF favors tumor progression (11,43). However, targeting PI3K or Pten (31), or Tgfbr1 and its signaling system (44) might be more effective.

**Figure 11:**
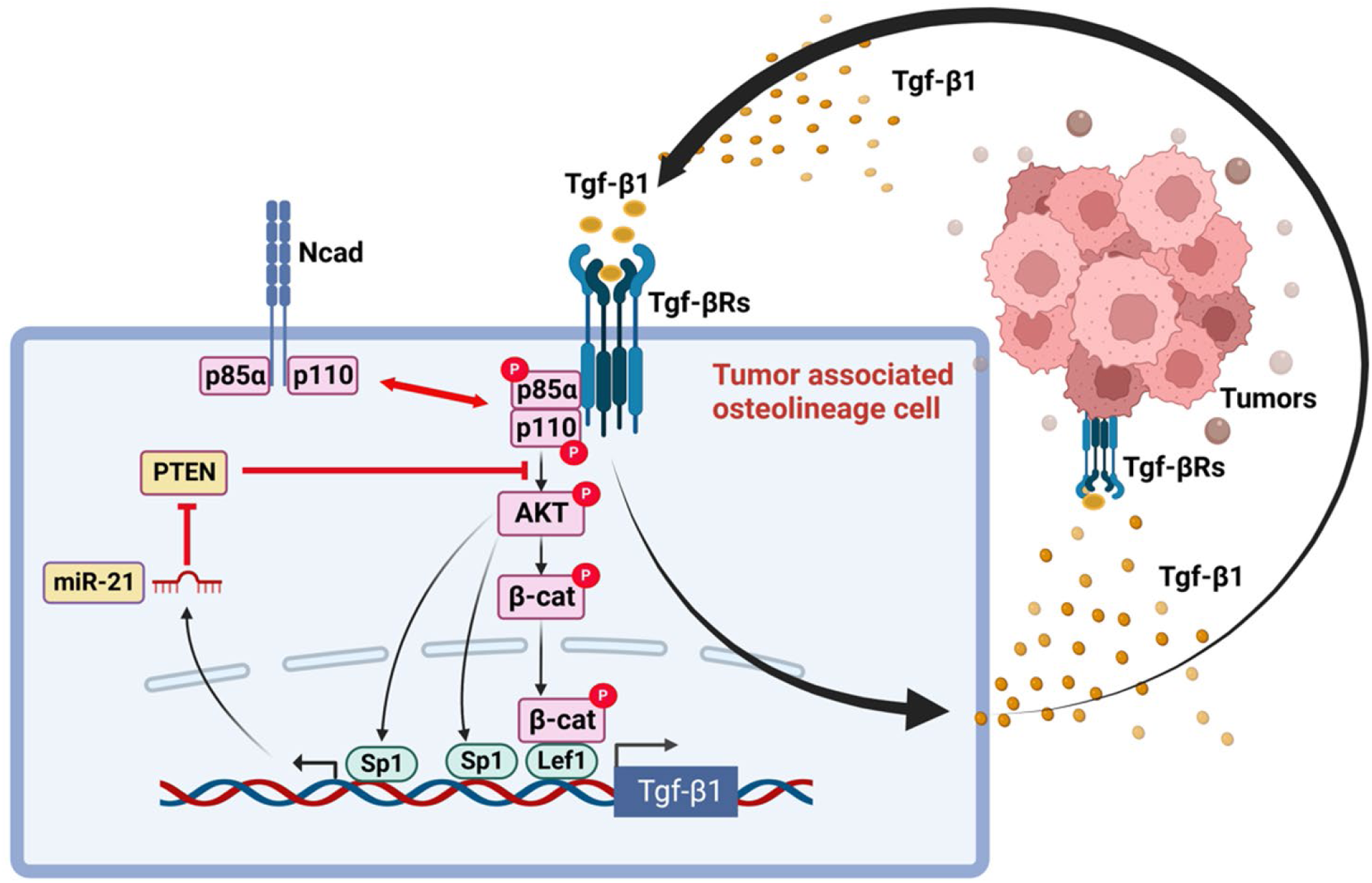
N-cadherin inhibits a pro-tumorigenic, Tgf-β1-driven, feed-forward autocrine loop in tumor associated osteolineage cells. We propose that Tgf-β1 in the TME stimulates tumor growth and activates tumor associated osteolineage cells *via* phosphorylation of two main PI3K components, p85 and p100, thus triggering a signaling cascade that results in AKT phosphorylation. In turn, activated p-AKT results in multiple downstream events, all leading to increased transcriptional activity of the *Tgfb1* promoter: 1) Sp1 recruitment to the promoter; 2) β-catenin stabilization by Ser552 phosphorylation, nuclear translocation, and promoter binding, most likely by complexing with Lef1; 3) increased miR-21 expression and post-transcriptional downregulation of the PI3K inhibitor Pten, enhancing this signaling cascade. This generates a positive feed-back loop that induces further autocrine Tgf-β1 production by TME osteolineage cells, fueling tumor growth. Ncad provides a break to this feed-forward “vicious” loop, by retaining the PI3K components, p85α and p110, away for signaling and thus, reducing PI3K signaling and Tgf-β1 production.

In conclusion, we discovered that by interfering with PI3K-Akt-β-catenin signaling in osteolineage cells, Ncad acts as a break to a Tgf-β1-driven positive feedback cycle that enhances autocrine production of Tgf-β1 and transcriptional up-regulation of the *Tgfb1* promoter. We propose that in tumor associated Osx+ cells, Ncad antagonizes their pro-tumorigenic action by inhibiting autocrine Tgf-β1 production. These findings add to our knowledge of how tumor cells interact with their microenvironment and disclose potential pharmacological targets.

## Acknowledgments

This work was supported by National Institute of Health grant R01 CA243383 and funds from the Barnes-Jewish Hospital Foundation (to RC), and by the Washington University Musculoskeletal Research Center (P30 AR074992).

## Significance

Tumor microenvironment cells with an osteogenic signature favor growth of primary mouse breast tumors. By interfering with PI3K-Akt-β-catenin signaling in these cells, the cell adhesion molecule Ncad inhibits a Tgf-β1-driven positive feedback cycle that enhances autocrine production of Tgf-β1 and tumor growth. These findings disclose potential new pharmacological targets to reduce tumor growth.

## References

1. de Visser KE, Joyce JA. The evolving tumor microenvironment: From cancer initiation to metastatic outgrowth. Cancer Cell 2023;41:374–403

2. Mrozik KM, Blaschuk OW, Cheong CM, Zannettino ACW, Vandyke K. N-cadherin in cancer metastasis, its emerging role in haematological malignancies and potential as a therapeutic target in cancer. BMC Cancer 2018;18:939

3. Tanaka H, Kono E, Tran CP, Miyazaki H, Yamashiro J, Shimomura T, et al. Monoclonal antibody targeting of N-cadherin inhibits prostate cancer growth, metastasis and castration resistance. Nat Med 2010;16:1414–20

4. Shintani Y, Fukumoto Y, Chaika N, Grandgenett PM, Hollingsworth MA, Wheelock MJ, et al. ADH-1 suppresses N-cadherin-dependent pancreatic cancer progression. Int J Cancer 2008;122:71–7

5. Augustine CK, Yoshimoto Y, Gupta M, Zipfel PA, Selim MA, Febbo P, et al. Targeting N-cadherin enhances antitumor activity of cytotoxic therapies in melanoma treatment. Cancer Res 2008;68:3777–84

6. Su Y, Li J, Shi C, Hruban RH, Radice GL. N-cadherin functions as a growth suppressor in a model of K-ras-induced PanIN. Oncogene 2016;35:3335–41

7. Turley RS, Tokuhisa Y, Toshimitsu H, Lidsky ME, Padussis JC, Fontanella A, et al. Targeting N-cadherin increases vascular permeability and differentially activates AKT in melanoma. Ann Surg 2015;261:368–77

8. Zhan DQ, Wei S, Liu C, Liang BY, Ji GB, Chen XP, et al. Reduced N-cadherin expression is associated with metastatic potential and poor surgical outcomes of hepatocellular carcinoma. Journal of Gastroenterology and Hepathology 2012;27:173–80

9. Yarom N, Stewart D, Malik R, Wells J, Avruch L, Jonker DJ. Phase I clinical trial of Exherin (ADH-1) in patients with advanced solid tumors. Curr Clin Pharmacol 2013;8:81–8

10. Beasley GM, Riboh JC, Augustine CK, Zager JS, Hochwald SN, Grobmyer SR, et al. Prospective multicenter phase II trial of systemic ADH-1 in combination with melphalan via isolated limb infusion in patients with advanced extremity melanoma. J Clin Oncol 2011;29:1210–5

11. Labernadie A, Kato T, Brugues A, Serra-Picamal X, Derzsi S, Arwert E, et al. A mechanically active heterotypic E-cadherin/N-cadherin adhesion enables fibroblasts to drive cancer cell invasion. Nat Cell Biol 2017;19:224–37

12. Wang H, Yu C, Gao X, Welte T, Muscarella AM, Tian L, et al. The osteogenic niche promotes early-stage bone colonization of disseminated breast cancer cells. Cancer Cell 2015;27:193–210

13. Fontana F, Xiang J, Su X, Tycksen E, Nassau R, Fox G, et al. N-cadherin in osteolineage cells modulates stromal support of tumor growth. Journal of Bone Oncology 2021;28:100356

14. Ricci B, Tycksen E, Celik H, Belle JI, Fontana F, Civitelli R, et al. Osterix-Cre marks distinct subsets of CD45-and CD45+ stromal populations in extra-skeletal tumors with pro-tumorigenic characteristics. Elife 2020;9:e54659

15. Hay E, Dieudonne FX, Saidak Z, Marty C, Brun J, Da Nascimento S, et al. N-cadherin/wnt interaction controls bone marrow mesenchymal cell fate and bone mass during aging. J Cell Physiol 2014;229:1765–75

16. Marie PJ, Hay E, Modrowski D, Revollo L, Mbalaviele G, Civitelli R. Cadherin-mediated cell-cell adhesion and signaling in the skeleton. Calcif Tissue Int 2014;94:46–54

17. Hay E, Nouraud A, Marie PJ. N-cadherin negatively regulates osteoblast proliferation and survival by antagonizing Wnt, ERK and PI3K/Akt signalling. PloS One 2009;4:e8284

18. Sudo H, Kodama HA, Amagai Y, Yamamoto S, Kasai S. In vitro differentiation and calcification in a new clonal osteogenic cell line derived from newborn mouse calvaria. J Cell Biol 1983;96:191–8

19. Su X, Esser AK, Amend SR, Xiang J, Xu Y, Ross MH, et al. Antagonizing Integrin beta3 Increases Immunosuppression in Cancer. Cancer Res 2016;76:3484–95

20. Chen JH, Shen C, Oh HR, Park JH. Glucocorticoids inhibit the maturation of committed osteoblasts via SOX2. J Mol Endocrinol 2022;68:195–207

21. Zhou S. TGF-beta regulates beta-catenin signaling and osteoblast differentiation in human mesenchymal stem cells. J Cell Biochem 2011;112:1651–60

22. Hamidi A, Song J, Thakur N, Itoh S, Marcusson A, Bergh A, et al. TGF-beta promotes PI3K-AKT signaling and prostate cancer cell migration through the TRAF6-mediated ubiquitylation of p85alpha. Sci Signal 2017;10

23. Weilbaecher KN, Guise TA, McCauley LK. Cancer to bone: a fatal attraction. Nat Rev Cancer 2011;11:411–25

24. Taurin S, Sandbo N, Qin Y, Browning D, Dulin NO. Phosphorylation of beta-catenin by cyclic AMP-dependent protein kinase. J Biol Chem 2006;281:9971–6

25. Fang D, Hawke D, Zheng Y, Xia Y, Meisenhelder J, Nika H, et al. Phosphorylation of beta-catenin by AKT promotes beta-catenin transcriptional activity. J Biol Chem 2007;282:11221–9

26. Chen XH, Lu LL, Ke HP, Liu ZC, Wang HF, Wei W, et al. The TGF-beta-induced up-regulation of NKG2DLs requires AKT/GSK-3beta-mediated stabilization of SP1. J Cell Mol Med 2017;21:860–70

27. Fang Q, Tian M, Wang F, Zhang Z, Du T, Wang W, et al. Amlodipine induces vasodilation via Akt2/Sp1-activated miR-21 in smooth muscle cells. Br J Pharmacol 2019;176:2306–20

28. Doumpas N, Lampart F, Robinson MD, Lentini A, Nestor CE, Cantu C, et al. TCF/LEF dependent and independent transcriptional regulation of Wnt/beta-catenin target genes. EMBO Journal 2019;38

29. Hay E, Buczkowski T, Marty C, Da Nascimento S, Sonnet P, Marie PJ. Peptide-based mediated disruption of N-cadherin-LRP5/6 interaction promotes Wnt signaling and bone formation. J Bone Miner Res 2012;27:1852–63

30. Revollo L, Kading J, Jeong SY, Li J, Salazar V, Mbalaviele G, et al. N-cadherin restrains PTH activation of Lrp6/beta-catenin signaling and osteoanabolic action. J Bone Miner Res 2015;30:274–85

31. Chalhoub N, Baker SJ. PTEN and the PI3-kinase pathway in cancer. Annu Rev Path 2009;4:127–50

32. Meng F, Henson R, Wehbe-Janek H, Ghoshal K, Jacob ST, Patel T. MicroRNA-21 regulates expression of the PTEN tumor suppressor gene in human hepatocellular cancer. Gastroenterology 2007;133:647–58

33. Massague J, Sheppard D. TGF-beta signaling in health and disease. Cell 2023;186:4007–37

34. Fontana F, Hickman-Brecks CL, Salazar VS, Revollo L, Abou-Ezzi G, Grimston SK, et al. N-cadherin Regulation of Bone Growth and Homeostasis Is Osteolineage Stage-Specific. J Bone Miner Res 2017;32:1332–42

35. Nagafuchi A. Molecular architecture of adherens junctions. Curr Opin Cell Biol 2001;13:600–3

36. Troyanovsky SM. Mechanism of cell-cell adhesion complex assembly. Curr Opin Cell Biol 1999;11:561–6

37. Bienz M, Clevers H. Linking colorectal cancer to Wnt signaling. Cell 2000;103:311–20

38. Brembeck FH, Schwarz-Romond T, Bakkers J, Wilhelm S, Hammerschmidt M, Birchmeier W. Essential role of BCL9-2 in the switch between beta-catenin’s adhesive and transcriptional functions. Genes Dev 2004;18:2225–30

39. Hay E, Laplantine E, Frain M, Geoffroy V, Muller R, Marie PJ. Permanent N-cadherin overexpression in pre-osteoblasts decreases osteoblast differentiation in vitro and bone mass in vivo by antogonizing Wnt signaling. J Bone Miner Res 2006;21:S21

40. Hay E, Laplantine E, Geoffroy V, Frain M, Kohler T, Muller R, et al. N-cadherin interacts with axin and LRP5 to negatively regulate Wnt/beta-catenin signaling, osteoblast function, and bone formation. Mol Cell Biol 2009;29:953–64

41. Gu J, Tamura M, Yamada KM. Tumor suppressor PTEN inhibits integrin-and growth factor-mediated mitogen-activated protein (MAP) kinase signaling pathways. J Cell Biol 1998;143:1375–83

42. Sizemore GM, Balakrishnan S, Hammer AM, Thies KA, Trimboli AJ, Wallace JA, et al. Stromal PTEN inhibits the expansion of mammary epithelial stem cells through Jagged-1. Oncogene 2017;36:2297–308

43. Li G, Satyamoorthy K, Herlyn M. N-cadherin-mediated intercellular interactions promote survival and migration of melanoma cells. Cancer Res 2001;61:3819–25

44. Kim BG, Malek E, Choi SH, Ignatz-Hoover JJ, Driscoll JJ. Novel therapies emerging in oncology to target the TGF-beta pathway. J Hematol Onc 2021;14:55

