## Supplemental Material for "N-cadherin in osteolineage cells restrains breast cancer cell growth via inhibition of a PI3K-dependent, Tgf-β1-driven feed-forward loop"

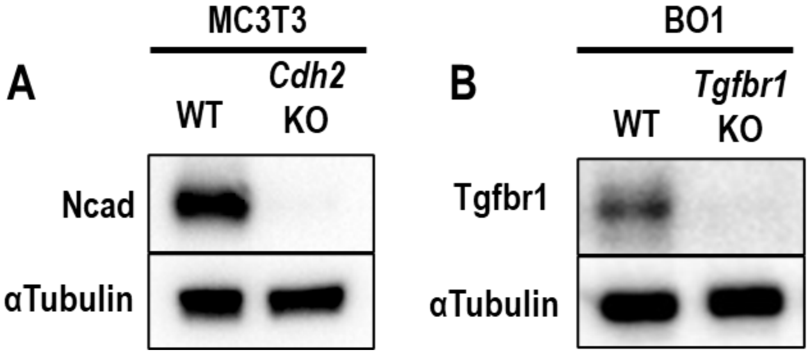


**Supplementary Figure 1. Ablation of *Cdh2* or *Tgfbr1* in MC3T3 or BO1 cells, respectively.** Western blot of whole cell lysates of (A) confluent WT or *Cdh2* KO MC3T3 cells using antibodies against N-cadherin (Ncad) or αTubulin; or (B) confluent BO1 breast cancer cells using antibodies against Tgfr1 or αTubulin.

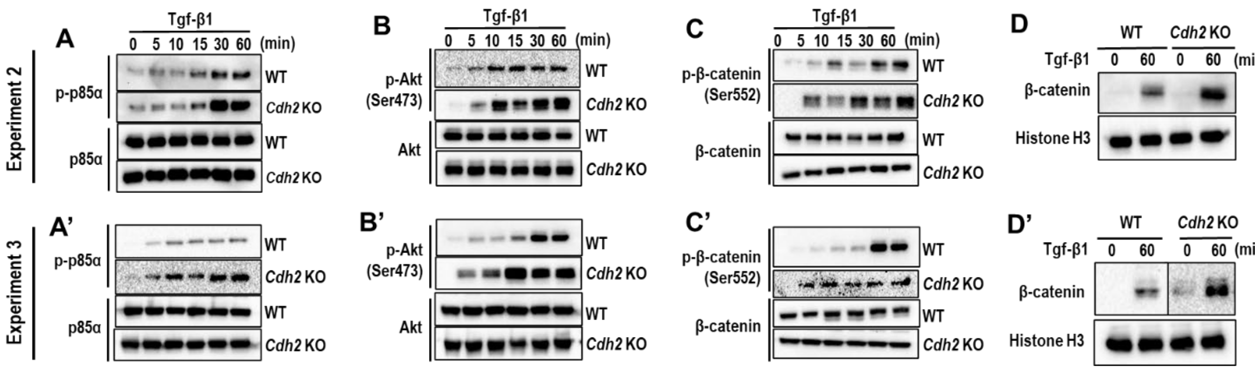


**Supplementary Figure 2. Hyperactivation of noncanonical Tgf-β1 signaling in *Cdh2* KO osteogenic cells.** Results of 2 additional, independent experiments identical to that shown in Fig. 1. Confluent MC3T3$^{WT}$ and MC3T3-E1$^{\mathit{Cdh2}KO}$ cells were exposed to Tgf-β1 (20 ng/ml) for different times, as indicated. Western blot of whole cell lysates using antibodies against (A, A’) p-PI3K p85α, PI3K p85α; (B, B’) p-Akt (Ser473), Akt, Ncad; and (C, C’) p-β-catenin (Ser552) or total β-catenin. (D, D') Nuclear extracts were obtained from confluent MC3T3$^{WT}$ cultures for Western blot analysis using antibodies against total β-catenin and histone H3 as loading control.

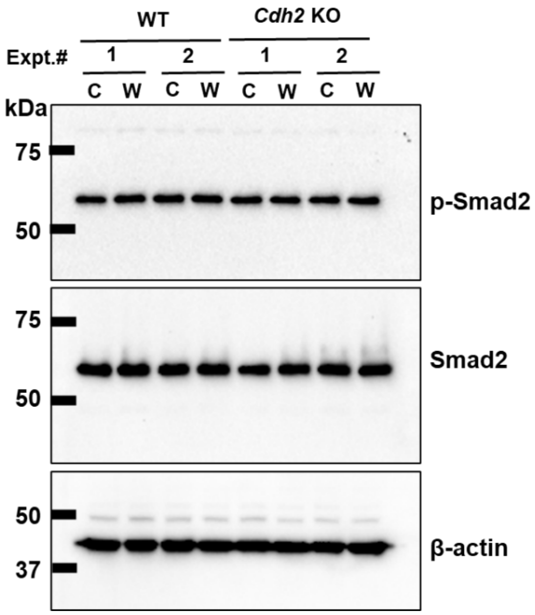


**Supplementary Figure 3. Unaltered activation of canonical Tgf-β1 signaling in *Cdh2* KO osteogenic cells upon PI3K inhibition.** Shown are the results of 2 independent experiments. Confluent MC3T3$^{WT}$ and MC3T3$^{Cdh2\ KO}$ cells were switched to αMEM without FBS and antibiotics for 6h, then incubated with the PI3K inhibitor, wortmannin (W: 1μM) or vehicle control (C) for 1 hour, followed by exposure to Tgf-β1 (20 ng/ml) for 1 hour. Western blot analysis of whole cell lysates incubated with anti-p-Smad2, anti-Smad2, or anti-β-actin antibodies.

**Supplemental Table 1**

List of Primers Used

| **For RT-qPCR** | Forward | Reverse |
|---|---|---|
| *Sp1* | CTCCAGACCATTAACCTCAGTGC | CACCACCAGATCCATGAAGACC |
| *Lef1* | ACTGTCAGGCGACACTTCCATG | GTGCTCCTGTTTGACCTGAGGT |
| *Tgfb1* | TGATACGCCTGAGTGGCTGTCT | CACAAGAGCAGTGAGCGCTGAA |
| *B2m* | ACAGTTCCACCCGCCTCACATT | TAGAAAGACCAGTCCTTGCTGAAG |
| *miR-21* | AGCTTATCAGACTGATGTTG | GAACATGTCTGCGTATCTC |
| *U6* | CTCGCTTCGGCAGCACAT | TTTGCGTGTCATCCTTGCG |
| *Tgfb1 promoter* | ATGGAGTGGAGTGTTGAGGG | CTTGCAGTCCATGGCATAGG |
| **For RT-PCR** | | |
| *Tgfb1 promoter* | CACGCAGATACCATCTACAGC | ACCCATGAGAAATACACGCTT |
